# Four Years of COVID-19: Saudi Arabia, Egypt and Pakistan Have the Highest Research Growth Rates From 2020-2023

**DOI:** 10.1101/2023.12.31.573759

**Authors:** Waseem Hassan, Mehreen Zafar

**Author notes:** Corresponding Author: Waseem Hassan, PhD.

## Abstract

We tried to assess the global research scholarly output after COVID-19 (from 2020 to 2023). Based on Scopus record, the world has produced 15, 041, 579 publications with 86, 165, 933 citations. We analyzed those countries, which have published at least 150, 000 research papers. For each country, we retrieved total number of publications, % growth rate, total citations, citations per paper, Field Weighted Citation Impact (FWCI), and % international collaboration. Twenty-seven (n=27) countries were found to be highly productive, with China leading the way in number of publications. Citation metrics are dominated by the USA, China, and European countries. Specifically, Switzerland, Netherlands, and Australia are notable for their high impact and influence. Saudi Arabia achieved the highest growth rate of 53.5%, and highest international collaboration (76.5%). Infact Saudi Arabia also attained high citations per article (8.8), and an FWCI of 1.63. While, Pakistan exhibited an 8.4 citations per article, FWCI of 1.54, growth rate of 34.9%, and collaborative percentage of 64.9%. Egypt also attained the 2^nd^ highest growth rate (n=36.1). Based on four (n=4) distinct performance metrics, Pakistan and Saudi Arabia were in the top ten group.

## 1.0 Introduction

Today (December 31, 2023), we will complete nearly four years, since the inception of the COVID-19 pandemic. This unprecedented global health crisis, has left an indelible mark on every facet of our lives. From the profound challenges faced by healthcare systems worldwide to the transformative shifts in how we work, learn, and interact, the journey through these four years has been one of resilience, adaptation, and collective efforts to overcome adversity. Bibliometric analysis (post-COVID-19) may provide valuable insights into the pandemic’s impact on global research output. It involves the quantitative analysis of publications, citations, and academic collaborations etc…

Recently Aksnes and Sivertsen published (1) a fascinating piece exploring the multifaceted dynamics of “Global trends in international research collaboration.” Using the Web of Science (WoS) Core Collection database, they analyzed 51.7 million papers with astounding 1.1 billion citations (from 1980 to 2021). They discussed in detail the regional differences and highlighted high-income nations as essential participants in the global research landscape (2, 3). The international collaboration exhibited a higher prevalence in smaller countries compared to their larger counterparts (4, 5). In another report, the role of shared investments in the European Union (EU) Framework programs, and correlation between financial commitments and a surge in collaborative activities among European countries is also presented (6).

The primary objective of the present study stems from a recognition of the transformative impact of the COVID-19 pandemic on global dynamics. Given the unprecedented changes, there arises a critical need to scrutinize and understand the evolving trends in international collaboration among countries. The timeframe of the study spans from 2020 to 2023 i.e. initiation and aftermath of the pandemic.

## 3.0 Materials and Methods

Scopus database, a robust and comprehensive platform renowned for its extensive coverage of scholarly literature, was employed. To present the research performance from 2020 to 2023, we employed various indicators. We focused on countries that exhibited a substantial research output i.e. at least 150,000 papers.

To gauge the productivity of the selected countries, we presented their overall % research growth rate. This metric serves as a key indicator of the expansion and dynamism for each country and may offer insights into the evolving nature of their scientific output.

To assess the quality of publications, we focused on total citations. By examining the cumulative citations received by each country’s publications, we aimed to capture the broader impact of their scientific contributions. We also presented the citations per paper (for each country), a metric designed to normalize the impact by considering the average number of citations each paper receives.

As part of our analysis, we introduced the concept of Field Weighted Citation Impact (FWCI). FWCI is a normalized indicator that accounts for the citation patterns within specific research fields. An FWCI greater than 1 indicates that the paper(s) has received more citations than the average for the field. In contrast, an FWCI less than 1 would imply that the paper(s) has received fewer citations than the average for the field. For instance, 1.20 means that a particular paper or set of papers has received 20% higher citations than the global average. We presented the FWCI for all selected countries.

Lastly, we examined the percentage of international collaboration for each country. This metric gauges the extent to which a country engages in collaborative research with international partners.

In summary, we employed a diverse set of indicators—number of publications, overall growth rate, total citations, citations per paper, FWCI, and the percentage of international collaboration—to offer a comprehensive and nuanced analysis of international research output among countries from 2020 to 2023.

## 4.0 Results and Discussion

From 2020 to 2023, the world has produced 15, 041, 579 publications with 86, 165, 933 citations. Twenty-seven countries have published at least 150,000 papers. Based on the number of publications, growth rate (%), total citations, citations per paper, Field Weighted Citation Impact (FWCI), and international collaboration (%), the top ten countries are presented in figure 1A-F.

**Figure 1A-F:**
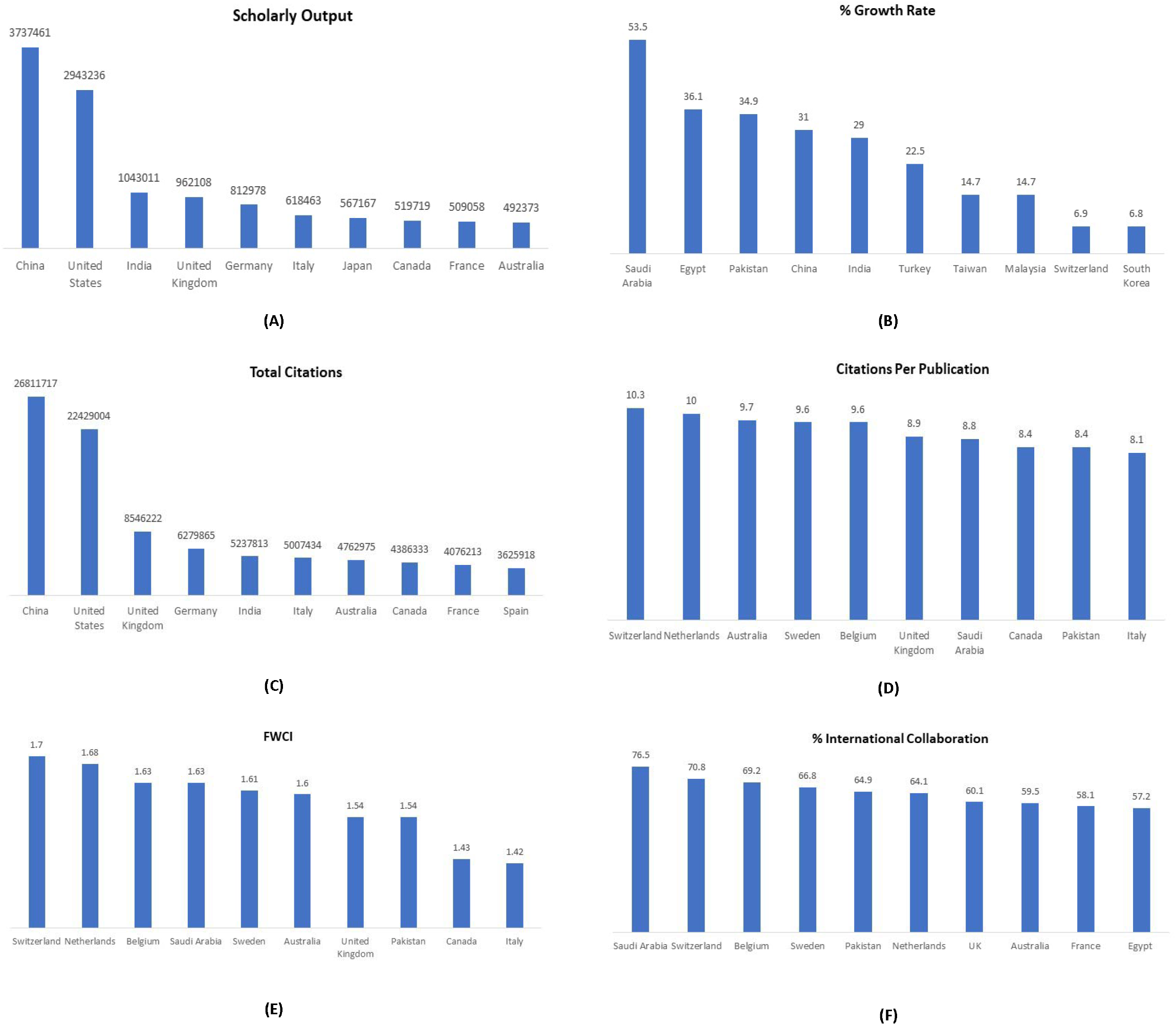
The top ten countries on the basis of scholarly output, % growth rate, total citations, citations per paper, FWCI and % international collaboration (from 2020-2023). The data was retrieved on December 28, 2023.

The data may offer a comprehensive snapshot of the scholarly landscape among various countries, highlighting their scholarly output, growth rates, citation metrics, and collaborative percentages.

For instance, China (n=3737461), United States (n= 2943236), and India (n= 1043011) are the powerhouses in terms of scholarly output, with China leading by a substantial margin.

Emerging economies such as Saudi Arabia (n= 53.5) and Egypt (n= 36.1), along with Pakistan (n= 34.9), are making substantial strides in increasing their scholarly output (as apparent from there %growth rates).

USA (n= 22429004) and China (n= 26811717), along with European nations, continue to be citation powerhouses, indicating the global impact of their research contributions.

Switzerland (n= 10.3), Netherlands (n= 10), and Australia (n=9.7) stand out in terms of citations per publication, suggesting high impact and influence.

Switzerland (n=1.7), Netherlands (n=1.68), and Belgium (n=1.63) also exhibited high Field-Weighted Citation Impact, emphasizing their impactful research across various fields. These countries stand out not just in terms of quantity but also the quality and influence of their research, as reflected in the field-weighted impact.

KSA (n=76.5), Switzerland (n=70.8), and Belgium (n=69.2) lead in the percentage of international collaboration, indicating a strong global research network.

Australia emerged as a standout contributor to the global scholarly landscape. Australia not only earned a position in the top 10 but has done so across a remarkable five key indicators, affirming its status as a powerhouse in the realm of academic research.

Canada, Italy, Switzerland, and the United Kingdom (UK) also demonstrated sustained excellence by securing positions in the top 10 across four key indicators. This impressive feat reaffirms their status as influential contributors to the world of academic research, showcasing consistent commitment to research output, impact, and collaboration.

Specifically, KSA exhibited a remarkable growth rate of 53.5%, placing it at the forefront of countries experiencing a surge in scholarly activity. This presents a significant commitment to expanding research endeavors. With an impressive score of 8.8 citations per publication, KSA demonstrated not only a quantitative increase in scholarly output but also the quality and impact of its research. KSA also holds a noteworthy FWCI of 1.63, showcasing that its research has an impact that exceeds the overall average. KSA also presented an exceptionally high international collaboration (n= 76.5%), underscoring its commitment to fostering global research networks. Similarly, the 2^nd^ highest growth rate was recorded for Egypt (from 2020-2023).

In the same vein, Pakistan demonstrated a substantial growth rate of 34.9%, reflecting a proactive approach to enhancing its scholarly output. With an average of 8.4 citations per publication, Pakistan not only increased research output but also the impact and recognition of its scholarly contributions on the global stage. Pakistan exhibited a notable FWCI of 1.54, indicating that its research has an impact that surpasses the average for its respective fields. With a collaborative percentage of 64.9%, Pakistan actively engaged in international research partnerships.

The present data highlights the positive trajectories of KSA and Pakistan in terms of scholarly growth, impact, and international collaboration, showcasing their evolving roles in the global research landscape. Similarly, a recent study (7) conducted in 2023 explored the productivity of authors in different countries. KSA and Pakistan were identified as having extremely productive authors, contributing significantly to the global scientific output. In another study (in 2018), the authors (8) reported the research growth rates of various countries. Notably, Egypt and Pakistan exhibited the highest growth rates in scientific publications during that specific year. We also reported the research growth rates of Pakistan in the fields of Chemistry (9) and Material Sciences (10), which revealed an unprecedented success, securing the top position among 40 and 50 countries, respectively.

However (in 2023), the researchers reported the scientific research landscape and some intriguing patterns in retraction rates (11). Surprisingly, both KSA and Pakistan exhibited high retraction rates compared to global averages. This raised significant questions about the quality control mechanisms, peer review processes, and research integrity within the scientific communities of these countries. The study prompted further investigations into the factors contributing to the observed retraction rates, such as potential issues with data validity, ethical concerns, or other systemic challenges. Combining critical measures to address challenges with strategic initiatives to support positive trends will contribute to a robust and reliable research environment in both countries. In summary, these studies may provide a nuanced view of the scientific research landscape, highlighting the complexities and variations within different countries. The findings not only contribute to the academic understanding of these regions but also offer opportunities for further investigation and improvement in the global scientific community.

## 5.0 Limitations

The present study has certain limitations.

The report provides descriptive data on growth rates, and international collaborations etc.. without delving into the underlying causes or reasons behind the observed trends. A more in-depth analysis is needed to uncover the drivers behind the high growth rates and collaborative efforts.

We just relied on Scopus and other databases were not analyzed.

The report lacks detailed information on the contextual factors influencing the research landscape in the studied countries. Understanding the socio-economic, political, and institutional contexts is crucial for interpreting the observed trends accurately.

The “Limited Exploration of Research Quality” highlights a significant gap in the present report, particularly in relation to assessing the original quality of research, with a specific focus on Pakistan. This limitation is crucial because understanding the quality of research output is integral to comprehensively evaluating a country’s scientific contributions.

Detailed insights into research budgets and expenditure were not provided. Understanding how countries allocate and utilize research funds is essential for evaluating the sustainability and effectiveness of research initiatives.

The report does not highlight specific areas of research or scientific advancements. Focusing on high-impact research areas, emerging technologies, and breakthroughs would provide a more forward-looking perspective on the global research landscape.

The report covers a specific timeframe (2020-2023) but does not conduct a longitudinal analysis to assess trends over an extended period. Longitudinal studies are essential for identifying patterns, shifts, and potential long-term impacts.

Descriptive data may lead to generalizations, and the report might not capture the diversity and nuances within each country’s research landscape. A more detailed study would involve a granular analysis of different research disciplines, institutions, and regions within each country.

While the report acknowledges international collaborations, it does not explore the nature and effectiveness of these collaborations. A detailed study could analyze collaborative models, success factors, and challenges faced in international research partnerships.

To address these limitations, future studies should incorporate detailed analyses, longitudinal perspectives, and in-depth explorations of critical aspects such as research quality, funding mechanisms, and the contextual factors shaping the research landscape in each country. Such studies would provide a more nuanced and comprehensive understanding of the dynamics within the global scientific community.

## Acknowledgment

None

